# wastewaterSPAdes: SARS-CoV-2 strain deconvolution using SPAdes toolkit

**DOI:** 10.1101/2022.12.08.519672

**Authors:** Anton Korobeynikov

## Abstract

**Motivation:** SARS-CoV-2 wastewater samples are extensively collected and studied because it allows quantitatively assess a viral load in surrounding populations. Additionally, SARS-CoV-2 strain deconvolution gives more insights into pandemic dynamics, and the uprising of new strains. Usually, the solution to the strain deconvolution problem starts with read alignment of wastewater short read sequencing data to the SARS-CoV-2 reference genome. After variants are called and their abundances are estimated, a reference database is used to assign variants to strains, select a subset of strains, and infer relative abundance of these strains based on some mathematical model. Assembly-based methods have its own strengths, but currently reside in the shadow of alignment-based methods.

**Results:** In this paper we propose a new assembly-based approach based on SPAdes toolkit codebase ––– wastewaterSPAdes, that is able to deconvolve wastewater without a need of read alignment. Our results show that watewaterSPAdes is able to accurately identify strains presented in a sample, and correctly estimate abundances for most of the samples.

**Availability:** https://cab.spbu.ru/software/wastewaterspades/

**Supplementary information:** Supplementary data are available at *Bioinformatics*

## 1 Introduction

Recent studies show that wastewater sampling is an effective way to predict local SARS-CoV-2 outbreaks, even before the number of confirmed cases starts to rise (Kitajima *et al*., 2020; Peccia *et al*., 2020). Wastewater samples usually contain multiple SARS-CoV-2 strains, and the problem of strain deconvolution naturally arises. Strain deconvolution is extremely useful as it allows researchers to track the dynamics of the spread of the virus in a community and monitor for potential outbreaks of novel strains. The the strain deconvolution problem can be split into two related subproblems: (i) identify the list of strains present in the sample, and (ii) assign each of the strains found its relative abundance. The solution to the problem is not straightforward, since different strains might have identical genome segments, as well as variants that can be shared between two or more strains. A few databases for distinctive variants of SARS-CoV-2 exist, the most notable are USHeR (Turakhia *et al*., 2021) and Pangolin (Rambaut *et al*., 2020). These databases can be used to determine a set of variants occurring in the sample. A few methods are being developed to solve the deconvolution problem, the most prominent of them is Frejya (Karthikeyan *et al*., 2022), which uses read alignment to call variations and use the USHeR database to deconvolve variants using the mixture model. All existing methods seem to use read alignment as a first step, while assembly-based methods are possible but sidelined. In this paper, we present an assembly-based method called wastewaterSPAdes.

## 2 Methods

In order to solve a SARS-CoV-2 decomposition problem, we first assemble sequencing data into an assembly graph using SPAdes (Prjibelski *et al*., 2020) but do not perform any extensive graph simplification procedures, except removing tips and bulges having *k*-mer coverage below 2. Then, every edge of the assembly graph with *k*-mer coverage above 10 is aligned into the SARS-CoV-2 reference genome together with its complement using the Smith-Waterman algorithm. Since the assembly graph contains both forward and reverse-complement sequences of the SARS-CoV-2 genome, we keep only one of two alignments with a better alignment score. The properties of the algorithm are chosen to keep alignments that have few small variations, even if they are close to the end of the aligned sequence, but to drop spurious alignments of reverse complements or contamination sequences. If the alignment has a variation, the variant type, sequence, and position are stored together with the coverage of the edge. Coverage of variants with the same position and sequence is accumulated. The assembly graph may have multiple occurrences of a single variant in the assembly graph, and some of them duplicate each other. In order to avoid excessive coverage accumulation, we only process variants that are located outside of the last *k* nucleotides of the edge, where *k* is the *k*-mer size used for assembly graph construction.

The next step is to determine the strains present in the sample based on collected variants. We use the USHeR database to identify a set of crucial variants for each strain. In order to select a subset of strains from the USHER database, for each strain we calculate the number of variants shared with the sample and the number of variants not present in the sample. Then we perform a multistep greedy procedure as follows.

- Sort all strains from the USHeR database in descending order based on the number of shared variants.
- Check stop condition, that the number of shared variants should be bigger than the number of missed variants, and the number of shared variants should be bigger than 3.
- Strain with the biggest number of shared variants is added to the list of strains.
- Shared variants for this strain are excluded in the next step of the algorithm.

Then for each strain from the list, we need to assign a probability, representing the relative abundance of the strain in the sample. In order to do that, for each strain we select a subset of variants that are not shared with other strains from the list. For each variation that is not shared, we calculate a relative abundance of that variant in the sample. So for each strain, we obtain a vector of numbers in (0,1] range. These numbers are averaged within the strain and normalized across the strains. The final number for each strain represents the abundance of the strain in the sample and this is the final result of the algorithm.

## 3 Results

We tested our approach on the synthetic data from (Karthikeyan *et al*., 2022). This dataset consists of 380 synthetic samples. For each sample, Aaron, Alpha, Beta, Delta, and Gamma SARS-CoV-2 strains were mixed in different known proportions ranging from 0% to 100%. These samples were sequenced using Covid-Seq and underwent an additional PCR amplification. As the results, these samples have extremely high and uneven coverage across the SARS-CoV-2 genome.

We assembled and estimated the strain abundances of each sample using wastewaterSPAdes. We compared these abundances against ground truth. In order to summarize the results, we classify the sample into the following groups and counted the number of samples falling into each group:

- *Correct*: 97 samples. All strains were identified correctly, and the difference between real and predicted abundances does not exceed 5%.
- *5-15% difference*: 179 samples. All strains were identified correctly, and the difference between real and predicted abundances does not exceed 15%, but at least one difference is above 5%.
- *More than 15% difference*: 94 samples. All strains were identified correctly, and at least one of the differences exceed 15%.
- *One lineage missing*: 0 samples. One of the strains was not identified.
- *Two or more lineage missing*: 0 samples. At least two strains were not identified.
- *One incorrect lineage*: 3 samples. One strain was incorrectly added to the list.
- *Two or more incorrect lineages*: 0 samples. Two or more strains were incorrectly added to the list.
- *Other*: 7 samples. All other cases. In our case, for these samples one strain was incorrectly replaced with other strain.

These results show that for most of the samples wastewaterSPAdes was able to correctly identify strains presented in the sample, and give reasonable estimates for the majority of the samples.

## 4 Conclusion

wastewaterSPAdes is the first assembly-based method capable of SARS-CoV-2 strain abundance estimation in wastewater samples. Potentially assembly-based methods are able to reduce biases in abundance calling. For such problems as strain deconvolution, reads are mapped to the reference genome without error correction procedures, therefore coverage estimates of each variation might be distorted. But in assembly-based methods, these errors are often removed from the graph as low-covered edges and don’t affect coverage calculations.

## Acknowledgements

The research was carried out in part by computational resources provided by the Resource Center “Computer Center of SPbU”. The authors are grateful to Saint Petersburg State University for the overall support of this work.

## References

Karthikeyan, S., Levy, J. I., De Hoff, P., Humphrey, G., Birmingham, A., Jepsen, K., Farmer, S., Tubb, H. M., Valles, T., Tribelhorn, C. E., et al. (2022). Wastewater sequencing reveals early cryptic sars-cov-2 variant transmission. Nature, 609(7925), 101–108.

Kitajima, M., Ahmed, W., Bibby, K., Carducci, A., Gerba, C. P., Hamilton, K. A., Haramoto, E., and Rose, J. B. (2020). Sars-cov-2 in wastewater: State of the knowledge and research needs. Science of The Total Environment, 739, 139076.

Peccia, J., Zulli, A., Brackney, D. E., Grubaugh, N. D., Kaplan, E. H., Casanovas-Massana, A., Ko, A. I., Malik, A. A., Wang, D., Wang, M., et al. (2020). Measurement of sars-cov-2 rna in wastewater tracks community infection dynamics. Nature biotechnology, 38(10), 1164– 1167.

Prjibelski, A., Antipov, D., Meleshko, D., Lapidus, A., and Korobeynikov, A. (2020). Using SPAdes de novo assembler. Current Protocols in Bioinformatics, 70(1).

Rambaut, A., Holmes, E. C., O’Toole, Á., Hill, V., McCrone, J. T., Ruis, C., du Plessis, L., and Pybus, O. G. (2020). A dynamic nomenclature proposal for sars-cov-2 lineages to assist genomic epidemiology. Nature microbiology, 5(11), 1403–1407.

Turakhia, Y., Thornlow, B., Hinrichs, A. S., De Maio, N., Gozashti, L., Lanfear, R., Haussler, D., and Corbett-Detig, R. (2021). Ultrafast sample placement on existing trees (usher) enables real-time phylogenetics for the sars-cov-2 pandemic. Nature Genetics, 53(6), 809–816.

